# Patterns of leaf, flower and fruit phenology and environmental relationships in a seasonal tropical forest in the Indian Eastern Himalaya

**DOI:** 10.1101/2024.09.29.615687

**Authors:** Aparajita Datta, Soumya Banerjee, Rohit Naniwadekar, Khem Thapa, Akanksha Rathore, Kumar Thapa, Turuk Brah, Tali Nabum, Ushma Shukla, Swati Sidhu, Noopur Borawake

**Affiliations:** Nature Conservation Foundation, 1311, “Amritha”, Vijayanagar 1st Stage, Mysuru, Karnataka, India 570017; BITS Pilani, Hyderabad Campus, Secunderabad, Telangana 500078

**Keywords:** Arunachal Pradesh, climate, cyclicity, photoperiod, seasonality, solar irradiance, temperature, tree phenology

## Abstract

Tree phenology plays an important role in determining the structure and function of tropical forest communities. However, there are few long-term studies on tree phenology from South Asia. We monitored 716 trees of 54 species monthly from 2011 to 2023 for leaf flush, flowering, and fruiting in Pakke Tiger Reserve, Arunachal Pradesh, India. We examined monthly patterns in the percentage of species and trees in leaf flush, flower and fruit and characterized phenological seasonality using circular statistics. Flowering periodicity was classified using Fourier analysis and we examined the relationships between phenological activity and temperature, rainfall, solar radiation and daylength using GAMLSS. Leaf flush and flowering were moderately seasonal, peaking in the warm dry season months of March to May. Fruiting patterns and their seasonality differed among dispersal modes. At the community level and for bird-dispersed species, fruiting was bimodal and relatively aseasonal, peaking in April and October. The highly seasonal fruiting of mammal-dispersed species peaked in October, while that of mechanically-dispersed species was bimodal and concentrated in the dry season months. The majority of species (78.13%) and trees (51.17%) flowered annually. Daylength, solar radiation and minimum temperature had significant nonlinear effects on phenology. This indicated the existence of narrow ranges of optimal climatic conditions for phenology, which could be affected by climate change. Our study emphasizes the need for long-term monitoring to rigorously quantify phenological patterns, particularly in the context of rapid global change.

## 1. INTRODUCTION

Phenology is the study of periodicity in the life-cycle events of plants and animals, which are influenced by seasonal variations in climatic factors such as temperature and precipitation. Seasonal patterns of leafing, flowering, and fruiting in forest trees have consequences for the life-history and breeding of a wide range of animal taxa.

The importance of environmental factors such as temperature, light, rainfall, relative humidity, solar radiation and edaphic factors on the vegetative and reproductive phenology of tropical trees have been emphasized in many studies in the last 30 years (Borchert 1983, Rathcke & Lacey 1985, Ashton *et al*. 1988, Kinnaird 1992, van Schaik *et al*. 1993, Tutin & Fernandez 1993, Chapman *et al*. 1999, Borchert *et al*. 2002, Kushwaha *et al*. 2011, Wright & Calderón 2018). Biotic factors may also play a role in driving flowering and fruiting (Rathcke & Lacey 1985, van Schaik *et al*. 1993). Trees may be adapted to aggregate or segregate flowering to ensure high pollination success or seed set (Rathcke & Lacey 1985, van Schaik *et al*. 1993, Bolmgren & Eriksson 2015) while fruiting patterns may be segregated (competition avoidance) or aggregated based on migrant and resident frugivore abundance (e.g. Kimura *et al*. 2001, Burns 2002). The major abiotic factors that have been identified as flowering cues are photoperiod, temperature, solar radiation and moisture (Rathcke & Lacey 1985, Chapman *et al*. 1999, Wright & Calderón 2018). While minimum temperature is an important proximate cue for flowering (Ashton *et al*. 1988, Tutin & Fernandez 1993), some earlier studies (Opler et al. 1976, Foster 1982a, Borchert 1983) have indicated that flowering is induced by rainfall.

Abiotic factors have been believed to be unimportant in stimulating fruit ripening (Rathcke & Lacey 1985), although moisture has been hypothesized to influence ripening secondarily by affecting fruit metabolism (Lacey 1980, Gautier-Hion 1990). Many studies have found that dehiscent dry fruits ripen in the dry season and fleshy fruits mature in the rainy season (Foster 1982a, Gautier-Hion 1990, Lieberman 1982, Chapman *et al*. 1999, Kitamura 2002, Cortes-Florés 2013).

It has been suggested that proximal environmental cues are dependent on whether water availability is a limiting factor (Graham *et al*. 2003, Zimmerman *et al*. 2007). Photoperiod and solar irradiance have been hypothesized to be the proximal climatic cues for the initiation of leaf flush and flowering at sites where water limitation is low, which maximizes photosynthetic efficiency (van Schaik *et al*. 1993, Wright 1996, Zimmerman *et al*. 2007).

The degree of climatic seasonality varies across tropical systems and is primarily influenced by the movement of the Inter-Tropical Convergence Zone (ITCZ) (Hastenrath 2012). Past studies have established that tropical forests in South-east Asia, Africa and the Neotropics are marked by seasonality (van Schaik *et al*. 1993, Sakai 2001). However, seasonality in phenological patterns may not always correspond to climatic seasonality. (Morellato *et al*. 2000, Sakai 2001).

Most studies on tropical forest phenology have been carried out in aseasonal forests near the equator, while there are fewer long-term studies in the more seasonal tropical forests, including tropical monsoon forests. There is limited information about long-term phenological patterns especially from India (latitudinal extent from 9 to 35° N) with very few long-term studies (see Ramaswami *et al*. 2019). Plant phenology studies from the seasonally dry tropical forests of India have been carried out primarily at sites dominated by dry deciduous and dry evergreen forests (e.g. Singh & Kushwaha 2006, Selwyn & Parthasarathy 2006, Kushwaha *et al*. 2011). The tropical forests in Arunachal Pradesh in the Eastern Himalaya at 27° North are among the world’s northernmost tropical rainforests (Proctor *et al*. 1998). Insights into the patterns of tree phenology from these more seasonal tropical forests in higher latitudes are limited (e.g. Shukla & Ramakrishna 1982, Kikim & Yadava 2001). Therefore, it is vital to relate plant phenological patterns in the northern tropical forests with those observed in other parts of the tropics and interpret regional variation in tree phenology in light of prevailing climate patterns. Establishing the relationship between seasonal climatic patterns and tree phenology is also vital for determining the nature of cyclicity of phenological patterns (Newstrom *et al*. 1994, Sakai 2001). In the Asian tropics, supra-annually occurring synchronous flowering has been documented extensively from dry dipterocarp-dominated and wet evergreen forests (Tsuji *et al*. 2023). However, plant reproductive cyclicity has been understudied in the more seasonally wet forest sites in tropical Asia, where annual cycles could be more common. Given the sensitivity of plant phenology to inter-annual variations in climatic conditions, it is especially important to have long-term data on plant phenology patterns to track possible effects of climatic fluctuations and environmental changes on tree species and forest ecosystems.

In this paper, we present patterns in vegetative and reproductive phenology of 54 tree species over 14 years. This is the only long-term study on tree phenology from this region. We examine the degree of seasonality and the relationship of key environmental variables to phenological patterns. We hypothesized that abundant soil moisture and its northerly location relative to other tropical forests would result in photoperiod, solar irradiance and temperature being the proximal cues for phenological activity. We also hypothesized that most species would exhibit annual reproductive cycles in response to predictably seasonal climate. Lastly, the observed patterns and the relationships were used to hypothesize and discuss the potential effects of climate change on tree phenology.

## 2. METHODS

### 2.1 Study Area

Pakke Tiger Reserve (862 km^2^, 92°36’ – 93°09’E and 26°54 – 27°16’N) is located in western Arunachal Pradesh, and is part of the Eastern Himalaya Biodiversity Hotspot. The elevation ranges from 200 – 1,700 m a.s.l. Pakke has a tropical climate. The long-term average annual rainfall is 2500 mm. The rainy season is from June to September (south-west monsoon) with some rainfall occurring due to the northeast monsoon (December to April). The cool dry winter are from November to February (Fig.1). Past temperature records (1983-1995) indicate that the mean (± SD) maximum temperature was 29.3°C ± 4.2 and the mean minimum temperature was 18.3° ± 4.7 (Datta 2001). For the period from 2011 to 2023, the mean maximum temperature is 30.5°C and the mean minimum temperature is 18.6°C.

**Fig. 1:**
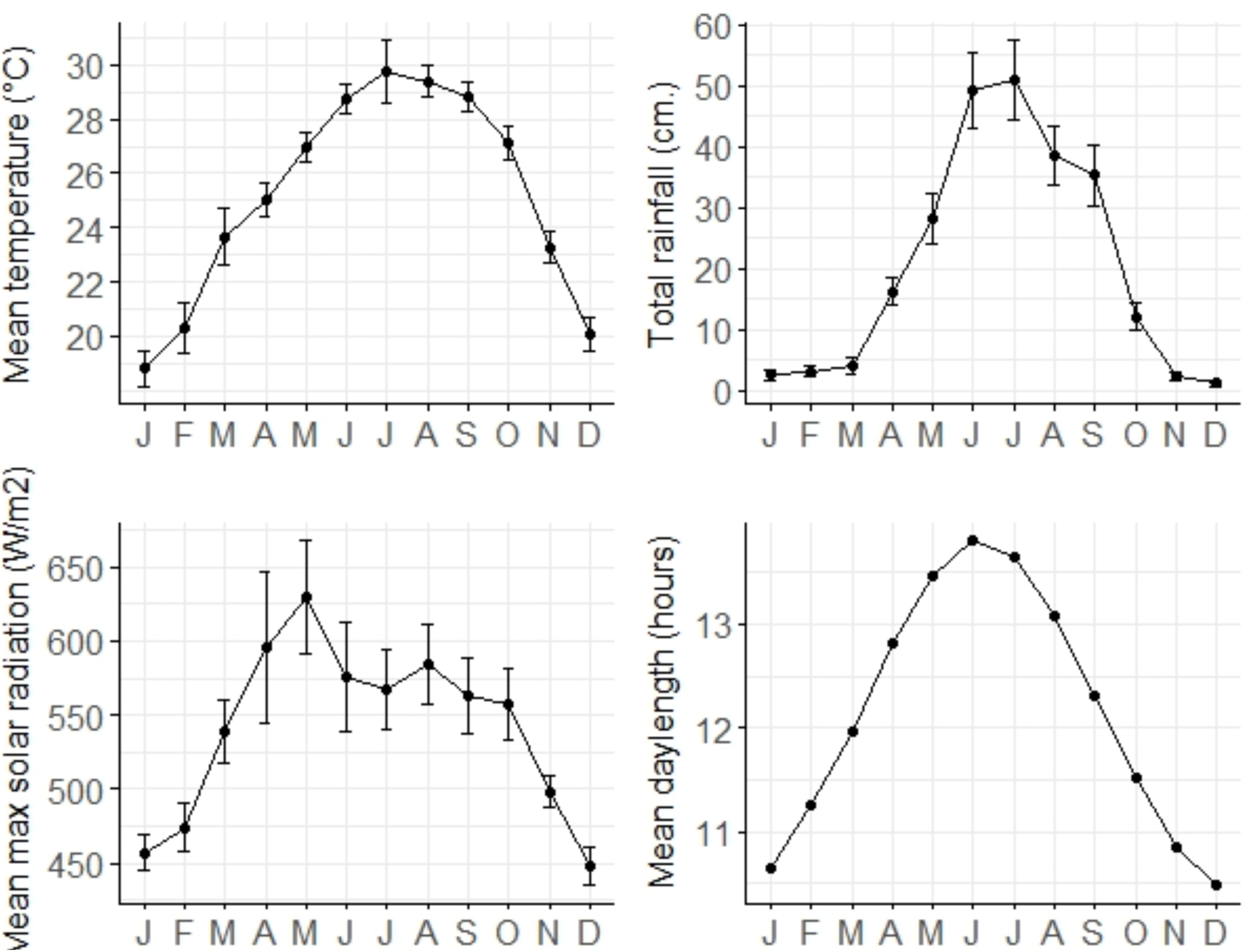
Monthly patterns in mean temperature, total rainfall, mean maximum solar radiation and daylength in the study area between January 2011 and February 2023. These patterns were based on data obtained from the weather station set up at Seijosa near Pakke.

The main forest type is Assam Valley tropical semi-evergreen forest 2B/C1 (Champion & Seth 1968). The forests are multi-storied and major plant families include Lauraceae, Euphorbiaceae, Myrtaceae and Meliaceae in the lower elevation forests (Datta 2001, Datta & Rawat 2008). Common tree species include *Monoon simiarum*, *Pterospermum acerifolium*, *Pterygota alata*, and *Duabanga grandiflora* (Page et al. 2022). Emergent species include *Tetrameles nudiflora*, *Ailanthus integrifolia* and *Liquidambar excelsa*.

### 2.2 Tree phenology monitoring

Tree phenology monitoring was initiated in January 2011. We monitored 716 reproductive trees of 54 selected tree species along 4 trails (Fig. S1) located in the lower elevation moist deciduous and semi-evergreen forests (150 - 600 m). The monitored trees were tagged/numbered with aluminium tags. Two to four trained observers using binoculars assessed the presence of each phenophases in the tree canopies. We recorded the presence/absence of flower buds, flowers, unripe fruit, ripe fruit, young leaf, mature leaf, senescent or old leaf or leafless and scored as 1 if any or all of these reproductive phases are present and as 0 if absent. This provides a total count of the number of trees of each species in various phenophases in every month. A particular tree may have young leaf, mature leaf, and senescent leaf at the same time. In such cases, all are recorded as being present on the tree. A few tree species are deciduous and are leafless for a certain period.

Till 2013, we had monitored the phenology only once a month (25^th^ – 28th of every month), the frequency was changed to fortnightly monitoring from 2014 (10^th^ –13^th^; and 25^th^ – 28^th^ of every month). For this paper, we have presented the results based on the monthly monitoring (25^th^ – 28^th^ of every month).

A prior vegetation and phenology study had been carried out (Datta 2001, Datta & Rawat 2008, Ramaswami *et al*. 2019) that had estimated average adult tree density in Pakke as 363 trees per ha (species richness: 162 species in 5.25 ha). The phenology tree sample represents 55% of this tree density with fifteen of the 20 top-ranked species represented. The tree species selected for phenology monitoring include eight mechanically-dispersed, nine mammal-dispersed, 21 bird-dispersed, and 16 dispersed by both birds and mammals based on Datta (2001) and Datta & Rawat (2008). Fifteen percent of the species monitored are wind-dispersed, while the rest are animal-dispersed. The sample size for 26 tree species was 20 - 25 individuals each, nine species had 10 - 19 individuals each, four species had 4 - 7 individuals each, and the remaining 14 species had < 3 individuals.

From 2011 to 2023, a total of 104 trees died (mean annual tree mortality of 1.27 %), of which 97 were replaced. The total number of trees monitored every month could vary based on the number of live trees in a month. The proportion or percentage of species and trees in each phenophase was calculated based on the total number of species and trees monitored every month. Phenological data was not available for July 2011, April and June-July 2015 and between March 2020 and April 2021 (during the Covid pandemic and due to lack of a research permit).

### 2.3 Weather monitoring

A weather station (H21-USB data logger) was set up in April 2011 in Seijosa, at the edge of Pakke (1 km straight-line distance from the reserve). The parameters recorded and downloaded from the weather station are temperature, rainfall, solar irradiance. Readings for these variables are recorded every hour, and to ensure that data is not lost due to any malfunctioning, we attempt to download and back-up the data every 15 days. However, we had complete weather data for all variables for 95 months during this period owing to occasional malfunctions.

### 2.4 Data Analysis

#### 2.4.1 Broad community-level phenological patterns

We estimated the mean monthly percentage of species and trees in each phenophase. These were depicted using line plots in order to visually represent phenological patterns over the course of the year. Patterns in fruiting phenology were examined at the community level and also separately for groups of species corresponding to the dominant seed dispersal modes: bird-dispersed, mammal-dispersed, animal-dispersed and mechanically-dispersed.

#### 2.4.2 Identifying cyclic patterns in reproductive (flowering) phenology

Fourier analysis was used to identify the predominant cycle of reproductive phenology at the level of the individual trees (Bush *et al*. 2016). The median flowering cycle length was estimated for 37 species for which at least 5 individual trees were sampled, using time series ranging from January 2011 to December 2019, in order to ensure that time series gaps were shorter than 3 months (Bush *et al*. 2016, Tsuji *et al*. 2023). Remaining gaps were filled using linear interpolation. A Daniell filter was used to smooth the raw periodogram in order to obtain a bandwidth of around 0.1 as per Bush *et al*. 2016. Flowering cycles were classified on the basis of their length as follows:- i) ≤10 months: sub-annual ii) 11-14 months: annual iii) ≥14 months: supra-annual. We did not estimate fruiting cycles as the high level of dioecy (up to 34% of monitored species) may have resulted in the overestimation of fruiting cycle length.

#### 2.4.3 Quantifying the degree of seasonality and mean and modal date of phenology

The degree of seasonality of various phenophases was quantified using circular statistics. For years in which all months were sampled (viz. 2012-2014, 2016-2019 and 2022), the monthly number of species or trees in each phenophase was considered to be circularly distributed, as per Hamann 2004 and Zimmerman *et al*. 2007. Rayleigh’s test was used to determine if phenological patterns were significantly seasonal (Jammalamadaka & Sengupta 2001, Morellato *et al*. 2010, Vogado *et al*. 2022). The length of the mean vector (denoted as r), which ranged from 0-1, was used as a measure of seasonality for sufficiently seasonal distributions (Morellato *et al*. 2010, Perina *et al*. 2019). The mean and the modal angle and the corresponding dates were also estimated using the number of species or trees sampled as weights. We examined for the existence of a linear trend in phenological activity by applying linear regressions on the mean phenological date converted into the corresponding day of the year. All analyses involving the use of circular statistics were carried out using the “circular” package in the R programming environment (Agostinelli & Lund 2017, R Core Team 2021).

#### 2.4.4. Modelling relationships between climate variables and phenology

We used the following climatic variables as covariates: i) mean maximum temperature ii) mean minimum temperature iii) total rainfall iv) proportion of rainy days v) mean of the daily maximum solar radiation and vi) mean daylength to determine the relationships between environmental variables and phenology on a monthly basis.

To accommodate complex, nonlinear relationships between environmental variables and phenology (Hudson *et al*. 2010, Sakai & Kitajima 2019, Vogado *et al*. 2022), we used the GAMLSS (Generalized Additive Models for Location, Scale and Shape) modelling framework (Rigby & Stasinopoulos 2005). Response variables for GAMLSS models were the proportion of trees in leaf flush, mature flower and ripe fruit in each month. The proportion of trees was preferred as the response variable in order to capture finer-scale community-level responses. A high degree of autocorrelation was observed at a 1 month lag for each phenological time series (r=0.32-0.80) (Fig. S2). Therefore, the value of the response variable in the previous month was used as a covariate (henceforth denoted as AR(1)), as per Hudson *et al*. 2010 and Vogado *et al*. 2022. Fruiting models at the dispersal-mode level exhibited poor fit, and hence, we present results for fruiting at only the community level.

We created models using additive functions of cubic smoothed splines corresponding to each covariate. The normal distribution was used to model the proportion of trees in each phenophase, as the beta distribution failed to converge for a number of flowering models. The “concurvity” metric was used to examine redundancy among splines of climate predictors used in the same model (Buja *et al*. 1989). Only those models with an estimated concurvity of <0.5 for smooth functions were retained in the final model set. We did not use more than 2 independent covariates per model in order to minimize overfitting. The Akaike Information Criterion (AIC) was used for model selection (Burnham & Anderson 2002) and conditional model averaging was used to derive estimates of beta coefficients for all models within <2 ΔAIC of the most parsimonious model. Modelling was carried out using the “gamlss” and “mgcv” packages in the R programming environment (Stasinopoulos *et al*. 2008, 2018, Wood 2017, R Core Team 2021).

## 3. RESULTS

### 3.1 Weather monitoring

Monthly patterns in temperature, precipitation and solar radiation appeared to show a high degree of seasonality. The highest mean monthly temperature (∼29°C) was observed in July, while the lowest mean monthly temperature (∼18°C) occurred in January. Monthly rainfall was highest during the south-west monsoon from June to September, (35 - 50 cm of monthly rainfall). Monthly rainfall was relatively low (≤5 cm) during the rest of the year. Solar radiation was highest in March-May, prior to the onset of the south-west monsoon, when cloudy skies resulted in a decrease in irradiance. Pakke had a comparatively high variation in daylength, ranging from 10.5 to 13.8 hours per month (Fig. 1).

### 3.2 Phenological patterns

Leaf flushing was comparatively lower in the cool dry season months of November to February in terms of both the percentage of species (23-36%) and the percentage of trees (6-14%). A significant increase in leaf flushing was observed at the commencement of the warm dry season in March and April. The percentage of species in leaf flush was highest in May (78.13%), while the percentage of trees in leaf flush was highest in April (53.50%) (Fig. 2).

**Fig. 2:**
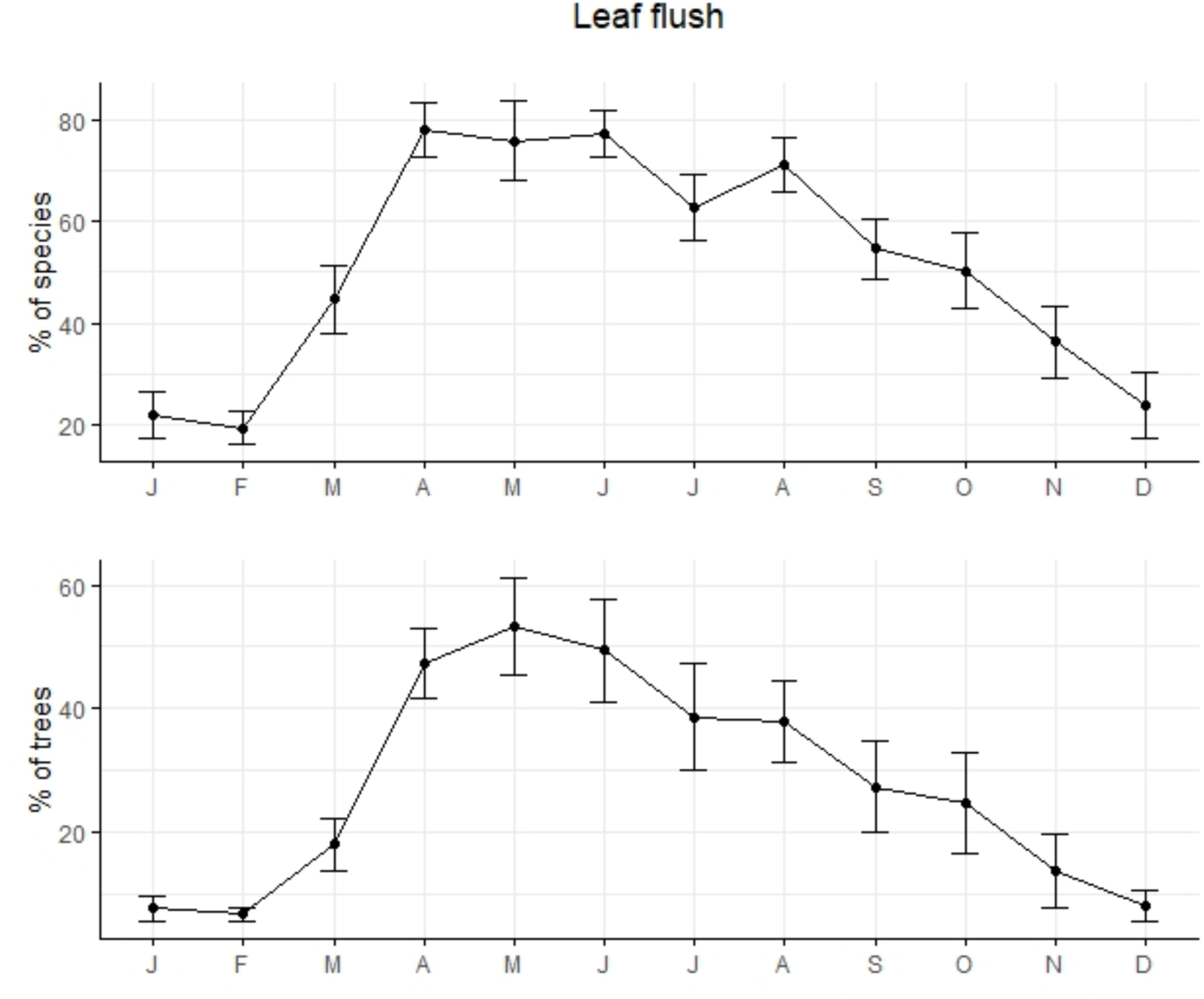
Monthly variation in percentage of species and trees in leaf flush at the community level in Pakke Tiger Reserve. Error bars represent the standard error of monthly values of percentage of species and trees in leaf flush. The above plot pertains to 54 species and 716 trees sampled between 2011 and 2023.

The occurrence of mature leaves was high throughout the year and was lowest in the warm dry season months of April and May. Leaf senescence peaked in the cool dry season months of November to February and declined thereafter, with the lowest values observed during the wet season in June.

Flowering commenced at the beginning of the warm dry season in March and April, and both the percentage of species and trees bearing flower buds were highest in April (35.24% and 16.14% respectively) (Fig. 3). The occurrence of mature flowers was fairly high between March and June (10.62-13.64% of trees in flower). There was a brief increase in floral activity at the end of the wet season in October, followed by low flowering in the cool dry months from November to February (Fig. 3). The warm dry season peak was more extended for mature flowers than it was for flower buds.

**Fig. 3:**
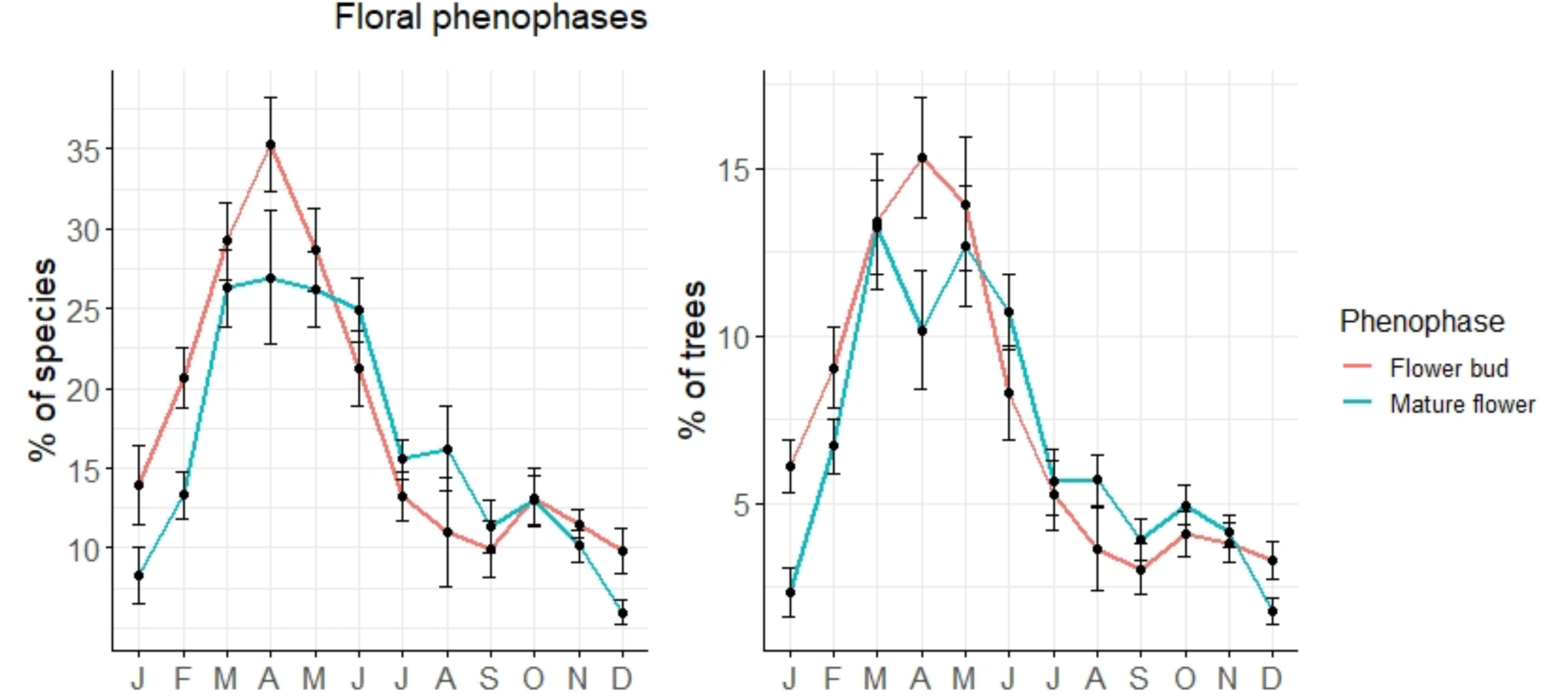
Monthly variation in percentage of species and trees in flower bud and mature flower at the community level in Pakke Tiger Reserve. Error bars represent the standard error of monthly values of the percentage of species and trees in the respective phenophases. The above plot pertains to 54 species and 716 trees sampled between 2011 and 2023.

Fruiting patterns varied across dispersal modes. The fruiting of animal-dispersed species appeared to be fairly high throughout the year, with the highest percentage of fruiting species being observed in April (20.05%) (Fig. 4). In terms of the percentage of trees, a bimodal pattern was observed, with the primary peak in April (8.25% of trees fruiting), while a smaller peak occurred in October. The fruiting of mechanically-dispersed species and trees was bimodal in nature and highest in the dry season months (Fig. 4).

**Fig. 4a:**
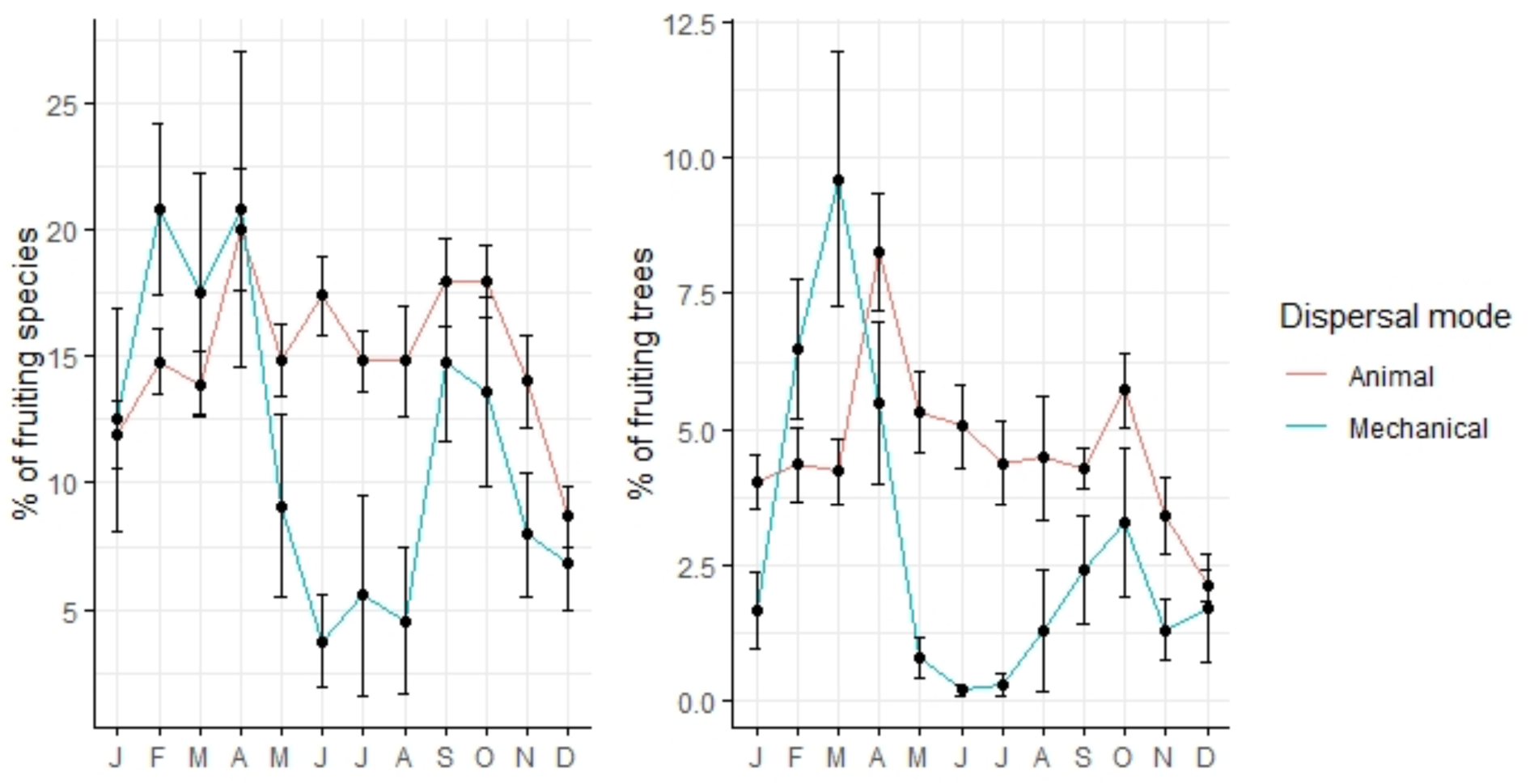
Monthly variation in the percentage of animal- and mechanically-dispersed species and trees in fruit in Pakke Tiger Reserve. Error bars represent the standard errors of the monthly values of the percentage of species in fruit. The above plot pertains to 565 animal-dispersed trees of 46 species and 151 mechanically-dispersed trees of 8 species.

**Fig. 4b.**
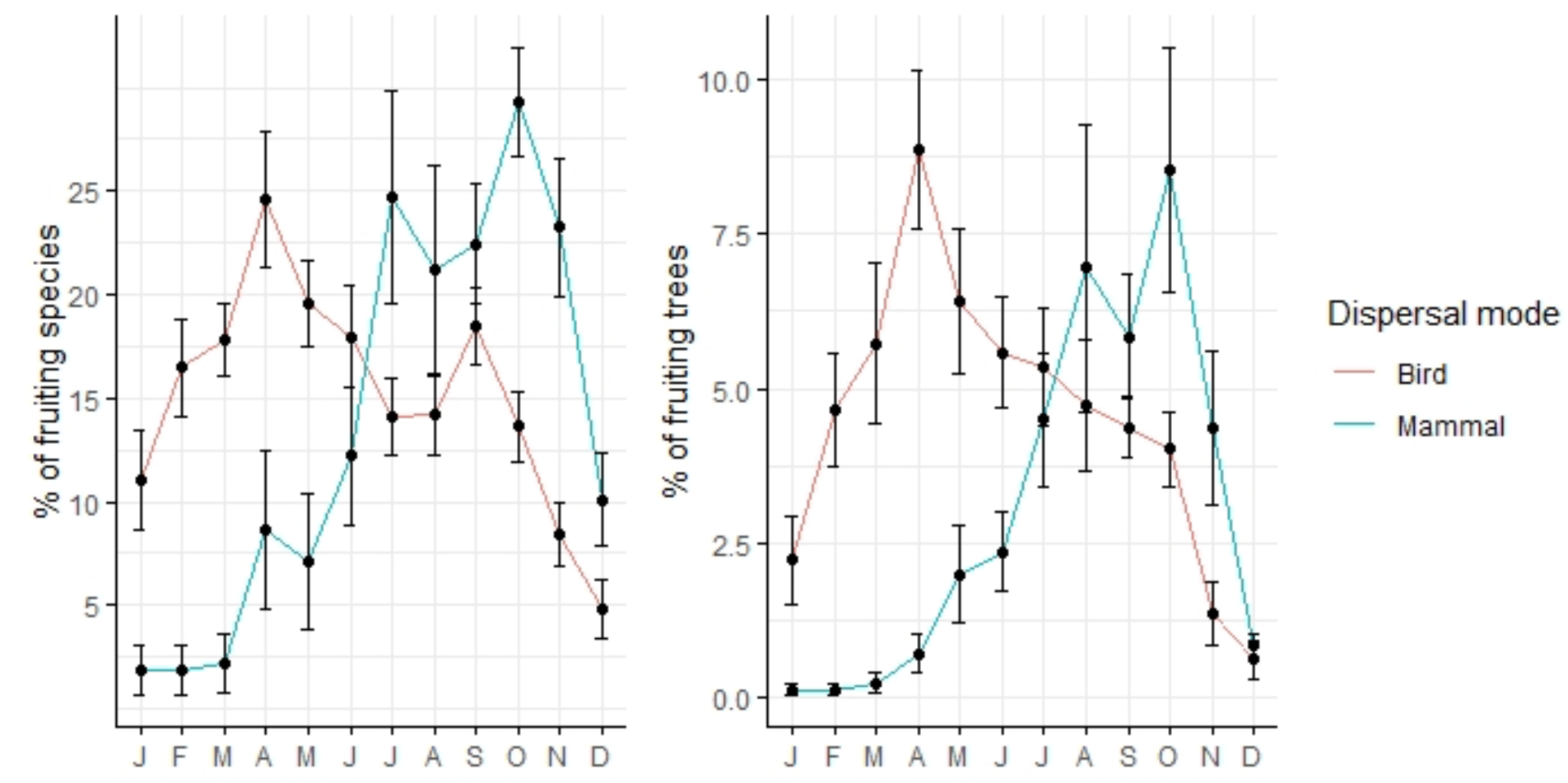
Monthly variation in the percentage of bird- and mammal-dispersed species and trees in fruit in Pakke Tiger Reserve. Error bars represent the standard errors of the monthly values of the percentage of species in fruit. The above plot pertains to 284 bird-dispersed trees of 21 species and 132 mammal-dispersed trees of 9 species.

Bird-dispersed species and trees also showed bimodal fruiting. The main peak was in April, when 24.6% of species and 8.86% of trees fruited. There was a smaller, secondary peak in October (Fig. 4). For mammal-dispersed species and trees, fruiting was relatively low in the warm dry season with its main peak shortly after the wet season in October, when 29.3% of species and 8.52% of trees were in fruit (Fig. 4). Fruiting of animal-dispersed species was generally low in the cool dry season.

### 3.3 Periodicity analysis

Fourier analysis was carried out for 643 trees belonging to 37 species. Flowering was acyclic for 98 trees (15.43% of the total sample). The majority of trees (329 or 51.17%) flowered annually, and sub- and supra-annual flowering was found in 117 (18.20%) and 99 (15.40%) trees respectively. Annual flowering occurred in 27 species, two species (*Prunus ceylanica* and *Alstonia scholaris*) flowered sub-annually while supra-annual patterns were observed in seven species (*Ailanthus integrifolia, Magnolia hodgsonii, Gynocardia odorata, Zanthoxylum rhetsa, Prasoxylon excelsum, Beilschmiedia assamica* and *Knema erratica*) (Table 1). The median length of the flowering cycle could not be estimated for *Artocarpus chaplasha* and the majority of individuals (14 out of 18) flowered acyclically.

**Table 1:** Median length of the flowering cycle in months for 36 species of trees in Pakke Tiger Reserve for which ≥5 individual trees were sampled and periodicity could be estimated using Fourier analysis. Species were categorized on the basis of the length of the flowering cycle as follows: i) ≤10 months: Sub-annual ii) 11-14 months: Annual iii) ≥15 months: Supra-annual.

| Species | Median flowering cycle length (months) | Periodicity classification |
| --- | --- | --- |
| *Dysoxylum gotadhora* | 12 | Annual |
| *Beilschmiedia assamica* | 15.43 | Supra-annual |
| *Dysoxylum cauliflorum* | 12 | Annual |
| *Prasoxylon excelsum* | 27 | Supra-annual |
| *Beilschmiedia sp.* | 10.8 | Annual |
| *Chisocheton cumingianus* | 12.75 | Annual |
| *Cryptocarya sp.* | 12 | Annual |
| *Actinodaphne obovata* | 12 | Annual |
| *Horsfieldia kingii* | 12 | Annual |
| *Knema angustifolia* | 19.5 | Supra-annual |
| *Zanthoxylum rhetsa* | 21.6 | Supra-annual |
| *Picrasma javanica* | 12 | Annual |
| *Cinnamomum cecicodaphne* | 12 | Annual |
| *Prunus ceylanica* | 6 | Sub-annual |
| *Baccaurea ramiflora* | 12 | Annual |
| *Gynocardia odorata* | 36 | Supra-annual |
| *Choerospondias axillaris* | 12 | Annual |
| *Endospermum chinense* | 12 | Annual |
| *Talauma hodgsonii* | 18 | Supra-annual |
| *Turpinia pomifera* | 12 | Annual |
| *Dillenia indica* | 12 | Annual |
| *Ailanthus grandis* | 21.2 | Supra-annual |
| *Alstonia scholaris* | 6 | Sub-annual |
| *Pterospermum acerifolium* | 12 | Annual |
| *Stereospermum chelonoides* | 12 | Annual |
| *Chukrasia tabularis* | 12 | Annual |
| *Tetrameles nudiflora* | 12 | Annual |
| *Bauhinia variegata* | 12 | Annual |
| *Sterculia alata* | 10.8 | Annual |
| *Polyalthia simiarum* | 12 | Annual |
| *Canarium resiniferum* | 12 | Annual |
| *Sterculia villosa* | 12 | Annual |
| *Livistona jenkinsiana* | 12 | Annual |
| *Elaeocarpus ganitrus* | 12 | Annual |
| *Aglaia spectabilis* | 12 | Annual |
| *Gmelina arborea* | 12 | Annual |

### 3.4 Patterns in seasonality and inter-annual variation

The monthly distributions of the number of species and trees in leaf flush and flower were seasonal in all years. Fruiting seasonality varied across dispersal modes. In terms of species in fruit, bird-dispersed species did not show seasonality in any of the years, while significant seasonality was not observed for mammal- and mechanically-dispersed species in 2 and 5 out of 8 years respectively (Table 2). All phenophases were significantly seasonal in terms of the number of trees in the respective phenophase. (Table 3).

**Table 2:** Circular statistical measures pertaining to the circular distribution of the number of species in each phenophase for each year in which all years were sampled (viz. 2012-2014, 2016-2019 and 2022). Distributions were weighted by the total number of sampled species for the estimation of mean angle. The mean angle was converted to the corresponding date. The Rayleigh’s test examines whether there is a statistically significant divergence from the uniform circular distribution and is used to examine whether the corresponding distribution was significantly seasonal. The length of the mean vector (denoted as R) is a measure of the degree of temporal aggregation of the circular distribution and is used as a measure of seasonality in this study. A statistically significant p value of the Rayleigh’s test is indicated by an asterisk (*). Where the Rayleigh’s test was not significant, values of statistical parameters are not presented.

| Year | Phenophase |  |  |  |  |  |  |
| --- | --- | --- | --- | --- | --- | --- | --- |
|  | Leaf flush | Flower bud | Mature flower | Ripe fruit (comm.) | Ripe fruit (bird) | Ripe fruit (mammal) | Ripe fruit (mech.) |
|  | Mean angle, date and R | Mean angle, date and R | Mean angle, date and R | Mean angle, date and R | Mean angle, date and R | Mean angle, date and R | Mean angle, date and R |
| 2012 | 186.34<br>(6<sup>th</sup> Jul) | 94.15<br>(4<sup>th</sup> Apr) | 144.27<br>(25<sup>th</sup> May) | NA | NA | NA | NA |
|  | R: 0.26* | R: 0.35* | R: 0.49* |  |  |  |  |
| 2013 | 169.00<br>(19<sup>th</sup> Jun) | 93.24<br>(3<sup>rd</sup> Apr) | 140.91<br>(22<sup>nd</sup> May) | NA | NA | 225.88<br>(16<sup>th</sup> Aug) | NA |
|  | R: 0.27* | R: 0.29* | R: 0.30* |  |  | R: 0.67* |  |
| 2014 | 175.02<br>(25<sup>th</sup> Jun) | 99.23<br>(10<sup>th</sup> Apr) | 113.16<br>(23<sup>rd</sup> Apr) | NA | NA | 260.10<br>(20<sup>th</sup> Sep) | NA |
|  | R: 0.37* | R: 0.27* | R: 0.25* |  |  | R: 0.55* |  |
| 2016 | 214.15<br>(4<sup>th</sup> Aug) | 103.17<br>(14<sup>th</sup> Apr) | 115.52<br>(26<sup>th</sup> Apr) | NA | NA | 248.26<br>(8<sup>th</sup> Aug) | NA |
|  | R: 0.32* | R: 0.32* | R: 0.39* |  |  | R: 0.53* |  |
| 2017 | 167.73<br>(18<sup>th</sup> Jun) | 90.44<br>(1<sup>st</sup> Apr) | 120.32<br>(1<sup>st</sup> May) | NA | NA | 238.37<br>(29<sup>th</sup> Aug) | 48.90<br>(18<sup>th</sup> Feb) |
|  | R: 0.34* | R: 0.28* | R: 0.27* |  |  | R: 0.53* | R: 0.57* |
| 2018 | 166.61<br>(16<sup>th</sup> Jun) | 144.18<br>(24<sup>th</sup> May) | 159.56<br>(10<sup>th</sup> Jun) | 152.73<br>(3<sup>rd</sup> Jun) | NA | 219.46<br>(10<sup>th</sup> Aug) | 87.92<br>(29 Mar) |
|  | R: 0.31* | R: 0.32* | R: 0.34* | R: 0.17* |  | R: 0.47* | R: 0.40* |
| 2019 | 183.85<br>(4<sup>th</sup> Jul) | 104.87<br>(15<sup>th</sup> Apr) | 117.74<br>(28<sup>th</sup> Apr) | NA | NA | 277.37<br>(8<sup>th</sup> Oct) | NA |
|  | R: 0.28* | R: 0.36* | R: 0.29* |  |  | R: 0.63* |  |
| 2022 | 155.87<br>(6<sup>th</sup> Jun) | 112.70<br>(23<sup>rd</sup> Apr) | 138.90<br>(20<sup>th</sup> May) | NA | NA | NA | NA |
|  | R: 0.34* | R: 0.43* | R: 0.34* |  |  |  |  |

comm. : community mech.: mechanically

**Table 3:**
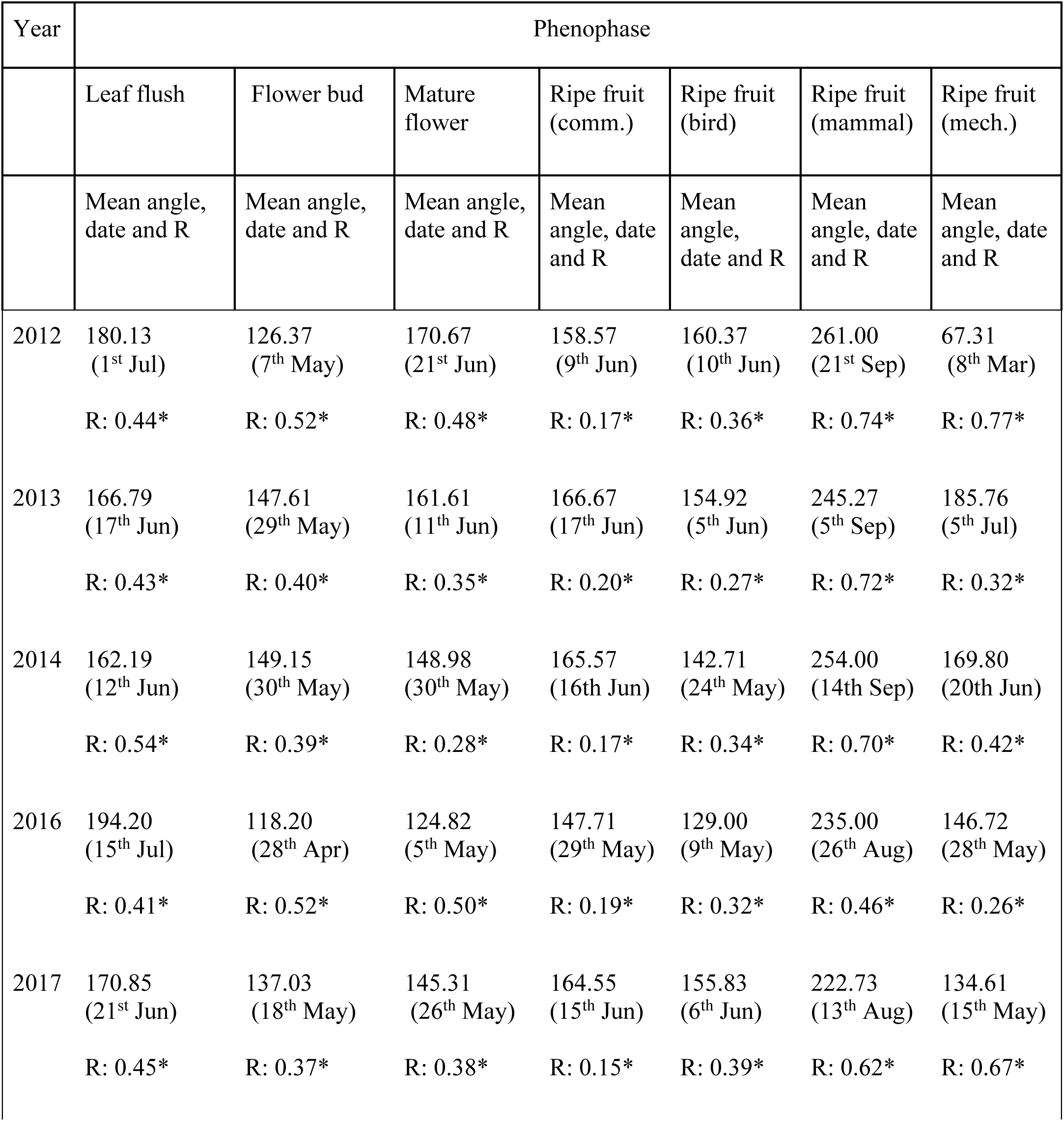
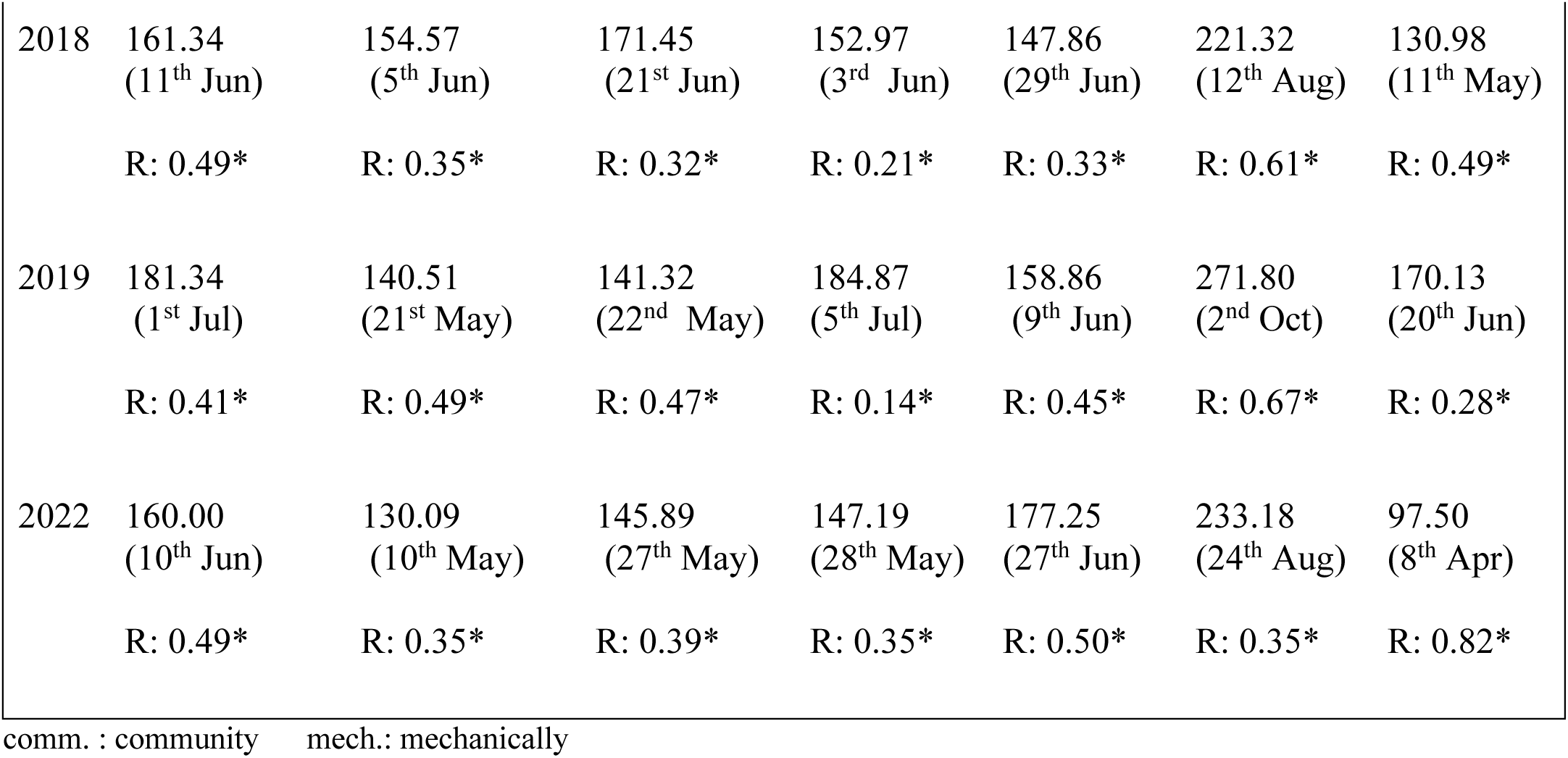
Circular statistical measures pertaining to the circular distribution of the number of trees in each phenophase for each year in which all years were sampled (viz. 2012-2014, 2016-2019 and 2022). Distributions were weighted by the total number of sampled trees for the estimation of mean angle.The mean angle was converted to the corresponding date. The Rayleigh’s test examines whether there is a statistically significant divergence from the uniform circular distribution and is used to examine whether the corresponding distribution was significantly seasonal. The length of the mean vector (denoted as R) is a measure of the degree of temporal aggregation of the circular distribution and is used as a measure of seasonality in this study. A statistically significant p value of the Rayleigh’s test is indicated by an asterisk (*). In cases where the Rayleigh’s test was not statistically significant, the mean angle, date and R value are not presented.

Overall, there was a statistically significant difference in the seasonality degree among the various phenophases in terms of the number of trees (ANOVA F = 9.751, df = 6, p<0.0001). We did not compare the degree of seasonality across phenophases in terms of the number of species, as not all phenophases were seasonal under this metric. The highest seasonality was shown by the fruiting of mammal-dispersed trees, followed by mechanically-dispersed trees and leaf flushing, with overall community-level fruiting showing the least seasonality (Table 3). Circular histograms indicated that the majority of monthly distributions were broadly unimodal or uniform, with some bimodality observed for the fruiting of bird-dispersed species in the year 2014 (Figs. S3 a) & b).

There was a significant degree of inter-annual variation in the timing of phenological activity. However, there was no linear trend in the mean date of occurrence of any of the phenophases (p=0.23-0.89).

The modal months of the various phenophases varied between years (Table S1). This was observed even in highly seasonal phenophases such as the fruiting of mammal-dispersed species. For instance, in 2017-2018 and 2022, the month with the highest percentage of mammal-dispersed trees in fruit was in the wet season (July, August and September respectively), while it occurred in October and November in other years (*viz.* 2012-2014, 2016, 2019) (Table S1).

### 3.5 Relationship between climate variables and phenology

We developed a total of 30; GAMLSS models incorporating additive combinations of only those predictors whose splines exhibited a low degree of concurvity. For all phenophases except for community-level fruiting, the autoregressive term AR (1) was statistically significant for all models (Table S3).

For leaf flush, models incorporating the effects of mean minimum temperature, solar radiation and daylength were best-supported (Table S2). Daylength had a statistically significant positive effect on leaf flushing (cs (daylength) = 0.048 ± 0.018, p<0.05) (Table S3).The smoothed functions of solar radiation and mean minimum temperature on leaf flushing had positive effects which were not, however, statistically significant (Table S3).

The most parsimonious model for flowering incorporated the additive effect of mean minimum temperature and solar radiation. Solar radiation had a positive effect (cs (solar radiation) = 0.012 ± 0.004, p<0.01) and minimum temperature had a negative effect on the occurrence of mature flowers (cs (minimum temperature) = -0.011 ± 0.005, p<0.05) (Table S3). For fruiting at the community level, models incorporating the effects of mean minimum temperature and solar radiation were found to receive the most support (Table S2). Solar radiation was found to have a statistically significant positive effect (cs (solar radiation) =0.006 ± 0.003, p<0.05). The overall effect of mean minimum temperature was not significant (Table S3).

Partial residual plots indicated substantial nonlinear effects of climate variables upon phenological activity. Leaf flushing increased almost monotonically with daylength till a maximum of around 13 hours of photoperiod, and declined thereafter (Fig. 5). Similarly, flowering declined sharply after minimum temperature reached 18°C and solar radiation of exceeded 600 W/m^2^ (Fig. 6). Fruiting was highest at a mean minimum temperature of 20°C, while the effect of solar radiation was nonlinear, with multiple inflection points (Fig. 7). Minimum temperature, solar radiation and daylength had the highest cumulative model support (Table S4).

**Fig. 5:**
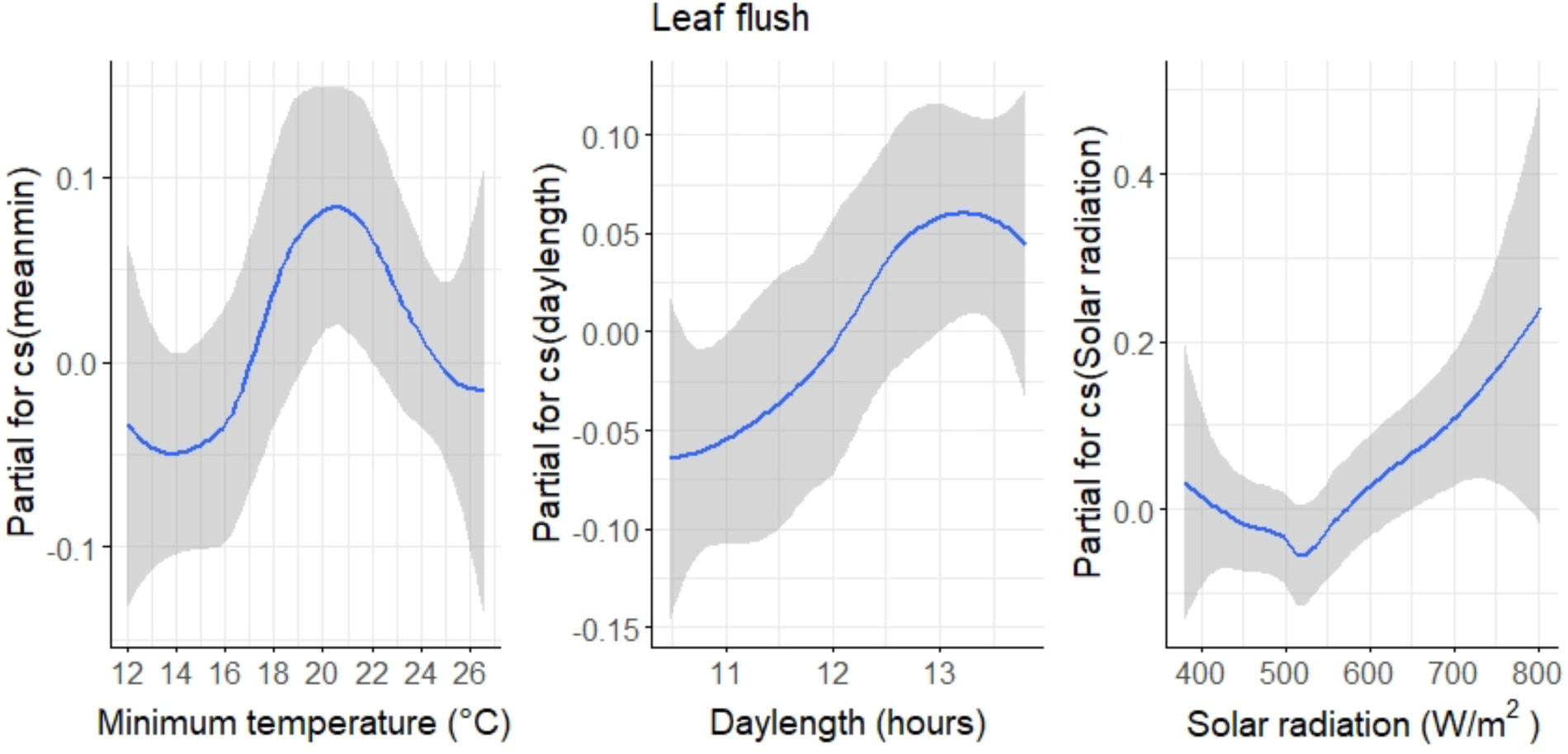
Partial residual plots depicting the relationship between cubic splines of mean minimum temperature, mean maximum solar radiation and the proportion of trees in leaf flush. The gray region depicts the 95% confidence intervals of the above relationships.

**Fig. 6:**
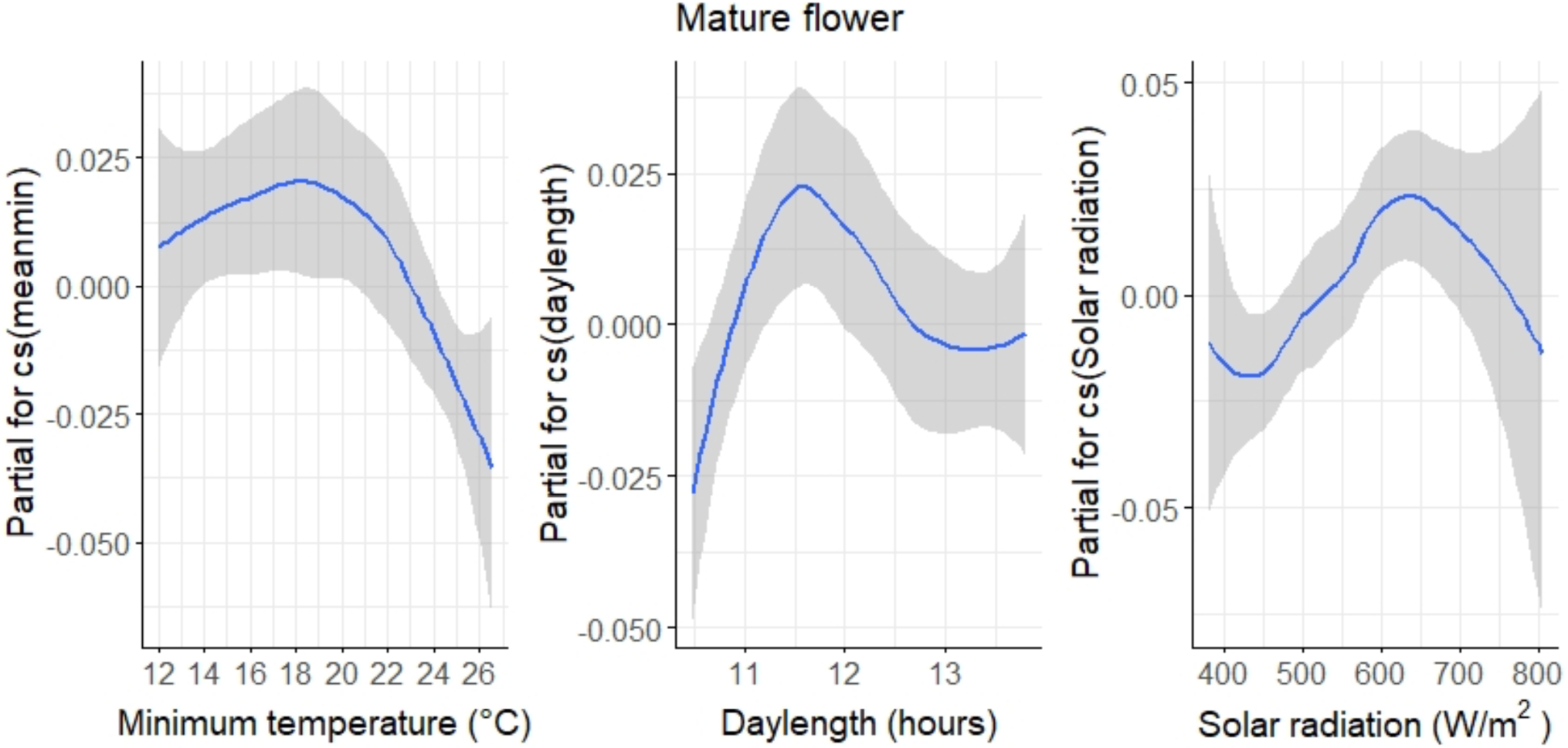
Partial residual plots depicting the effect of the cubic smoothed splines of mean minimum temperature and mean maximum solar irradiance (on the log scale) on the proportion of trees bearing mature flowers. The gray region depicts the 95% confidence intervals of the above relationships.

**Fig. 7:**
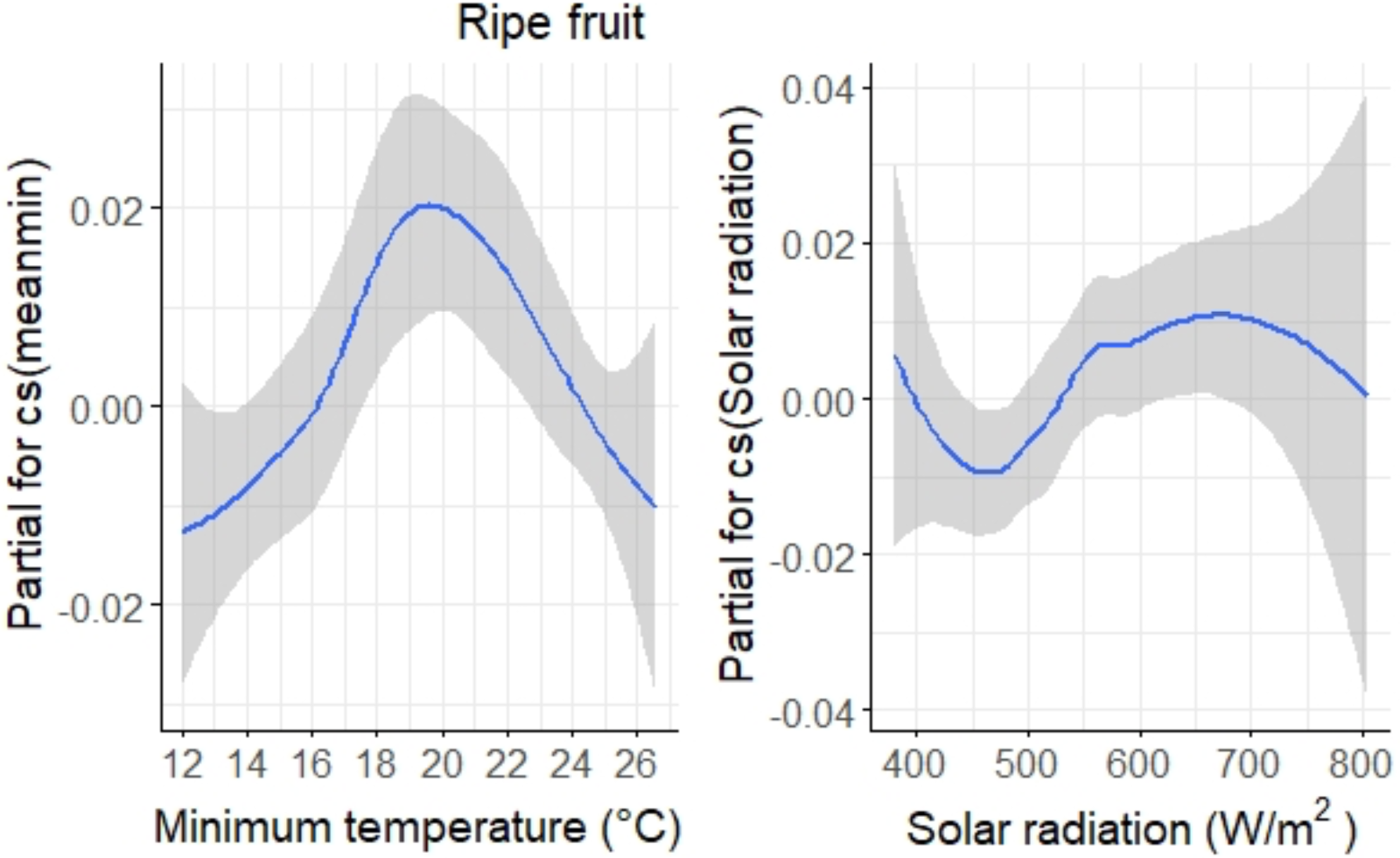
Partial residual plots depicting the effect of the cubic smoothed splines of mean minimum temperature and mean maximum solar irradiance (on the log scale) on the proportion of trees bearing ripe fruit. The gray region depicts the 95% confidence intervals of the above relationships.

## 4. DISCUSSION

### 4.1 Patterns of seasonality among phenophases

Our study indicated that broad community-level patterns of leaf flushing and flowering were moderately seasonal (r = 0.4-0.5), with varying levels of seasonality across phenophases and years. Leaf flushing and flowering activity increased from the annual low in the cool winter months and peaked in the warm dry season between March and May, however, it remained high into the beginning of the wet season in June.

Fruiting patterns and their degree of seasonality varied across seed dispersal modes and the metric under consideration (i.e. number of species versus trees in fruit). The fruiting of mammal-dispersed trees was more seasonal compared to that of bird- or mechanically-dispersed trees, and was the highest among all phenophases. Phenological patterns were broadly similar in terms of both analysed measures, viz. the percentage of species and trees in each phenophase. However, all phenophases showed higher seasonality with the percentage of trees. Although many studies have described community-level phenological patterns using only the proportion of species (e.g. Sun *et al*. 1996, Morellato *et al*. 2000, Selwyn & Parthasarathy 2006), this may underestimate the degree of phenological seasonality. This is particularly applicable where a relatively large number of trees are being monitored per species.

### 4.2 Phenological patterns and their relationships with climatic variables

Leaf flushing commenced in the warm dry season, similar to broader patterns of flushing observed across a precipitation gradient in tropical Asian forests (Shukla & Ramakrishnan 1982, Singh & Kushwaha 2006, Elliot *et al*. 2006, Hanya *et al*. 2013). In Pakke, leaf flush continued into the wet season, and the mean date of leaf flushing was in June and July in all years. However, the highest percentage of trees in flush were in the dry season (April-May).

Although proximate factors driving vegetative phenology are related to the tree water balance and its implications for water stress (Singh & Kushwaha 2005, Elliot *et al*. 2006), increasing daylength has been hypothesized to be a reliable proximal cue for leaf flushing in tropical Asia (Elliot *et al*. 2006, Kushwaha *et al*. 2011). This was validated by the statistically significant positive effect of photoperiod obtained through GAMLSS modelling in Pakke.

The concentration of flowering in the warm dry season months (March-April) in Pakke has been reported across seasonally dry tropical Asian forests (Singh & Kushwaha 2005, Mohandass *et al*. 2018). We observed a smaller peak in the post-monsoon dry season, which is when flowering was highest in the Afrotropics (Chapman *et al*. 1999, Babweteera *et al*. 2018). Both leaf flushing and flowering peaked in May, and this synchrony between the two phenophases at the community level has been widely reported across the tropics (Murali & Sukumar 1994, Kushwaha *et al*. 2011). Solar irradiance had a positive effect on flowering in Pakke, which has been reported from seasonally dry tropical forests (Hamann 2004, Zimmerman *et al*. 2007, Wright & Calderón 2018).

The negative effect of minimum temperature on the flowering of trees in Pakke is in conformity with similar effects reported for *Eucalyptus* spp. in tropical Australia (Hudson *et al*. 2010) and the different phenomenon of general flowering in aseasonal forests of South-east Asia (Chen *et al*. 2018, Ushio *et al*. 2020). However, other studies from the tropics have hypothesized a positive effect of temperature on flowering (Singh & Kushwaha 2005, Pau *et al*. 2018). Partial residual plots indicated that the negative effect of mean minimum temperature in Pakke was prominent only above a temperature of 20°C, while at lower temperatures, the relationship between minimum temperature and flowering appeared to be weakly positive. Thus, lower temperatures may have been a limiting effect restricting flowering.

Flowering cycles were predominantly annual at both the species and the individual level. The proportion of monitored trees showing annual patterns was higher in Pakke than that observed across 11 sites in tropical Africa (46%) (Adamescu *et al*. 2018), where annual patterns were also dominant.

Our study is among the first to rigorously quantify flowering cyclicity from a seasonally dry site in the Asian tropics, with previous studies having been carried out in aseasonal forests. (e.g. Sakai et al. 1999, Tsuji et al. 2023). The dominance of annual flowering in Pakke lends support to the importance of climatic seasonality on plant reproductive cyclicity (Newstrom et al. 1994, Sakai 2001, Kushwaha *et al*. 2011).

The differing monthly peaks of fruiting among the various seed dispersal modes resulted in an overall uniform fruiting at the community level. Differing fruiting patterns among dispersal syndromes in Pakke could be related to selective pressures related to reproductive success (Gautier-Hion 1990, Bolmgren & Eriksson 2005). The peak fruiting of bird-dispersed species in April currently coincides with the commencement of breeding season of most resident bird species, including frugivores. There appeared to be an advancement in the peak fruiting of bird-dispersed species from the months of June-July in the period 1997-2000 (Datta 2001, Ramaswami *et al*. 2019), although it is unclear whether this relates to climate change. Similarly, the dry season fruiting peak of mechanically-dispersed species favors fruit dehiscence and higher dispersal success due to greater wind speeds in the dry season (van Schaik *et al*. 1993, Griz & Machado 2001).

Solar irradiance had an overall positive effect on fruiting, which has also been reported from several tropical forest sites (Wright & Calderón 2006, Zimmerman *et al*. 2007, Chapman *et al*. 2018). The overall effect of mean minimum temperature on fruiting was not significant. However, fruiting increased with temperature up to a value of 20°C; similar nonlinear relationships between temperature and fruiting have been hypothesized in tropical Asia (Tutin & Fernandez 1993).

### 4.3 Relative importance of climate variables

Solar irradiance, temperature and photoperiod had a bigger effect on phenological patterns compared to precipitation-related variables. This supports the hypothesis that phenological patterns are driven more by light availability through solar irradiance and photoperiod in sites where soil moisture is plentiful (Wright 1996, Graham *et al*. 2003, Elliot *et al*. 2006, Zimmerman *et al*. 2007). In Pakke, with its high annual rainfall and northerly location, solar irradiance and photoperiod are likely to impose greater constraints on phenological activity through photosynthetic efficiency compared to rainfall. Minimum temperature is relatively low in Pakke (annual mean minimum of ∼12°C) compared to other lowland forests in tropical Asia. Therefore, low minimum temperatures may be a limiting factor on tree phenology. For most phenophases, minimum temperature received greater cumulative model support compared to irradiance and photoperiod. However, we suggest that the more intuitive effect on solar irradiance on photosynthetic efficiency would probably result in solar irradiance being the climate variable with the most significant ecological effect on phenology. The importance of solar radiation is further supported by its high degree of model support for all phenophases, whereas minimum temperature did not have a significant relationship with leaf flush or fruiting. Although rainfall did not appear to be a proximal cue of phenological activity, it may facilitate leaf expansion or seed dispersal through germination (Kushwaha *et al*. 2011, de Camargo *et al*. 2018).

Partial residual plots indicated that tree phenology was dependent on narrow optimal ranges of key climate variables. Therefore, anthropogenic climate change-mediated effects on climate variables (Parmesan 2007) relative to conditions for optimality could affect tree phenology. This could have adverse effects upon mutualistic relationships such as pollination or seed dispersal (Renner & Zohner 2018, Gérard *et al*. 2020). Our results highlight the need for robust modelling techniques such GAMLSS to account for the complex nonlinear relationships between climate variables and tree phenology.

### 4.4 Inter-annual variability in phenological patterns

We found substantial inter-annual variation in the timing and degree of seasonality of phenophases. For example, the fruiting of mechanically-dispersed species was seasonal only in 2017 and 2018 and for mammal-dispersed species in all years except 2012 and 2022. On the other hand, leaf flush and flowering was seasonal in all years. The modal months for each phenophase also varied across years.

Significant inter-annual variation in the amount and timing of flowering and fruiting has been reported from several other tropical forest sites (Polansky & Boesch 2013, Mendoza *et al*. 2018). Periodic climatic phenomena such as ENSO (El Nino-Southern Oscillation) and IOD (Indian Ocean Dipole) are known to influence long-term patterns of reproductive phenology in tropical forests (Chapman *et al*. 2018, Diem *et al*. 2018, Vogado *et al*. 2022), and their role in inter-annual variation in the phenological timing in Pakke needs to be examined. Species-specific phenological responses to climate change also need to be studied (Miller-Rushing *et al*. 2008, Calinger *et al*. 2013). Our study highlights the importance of climatic seasonality in determining phenological patterns in Pakke. It also serves as a crucial benchmark of knowledge on tree phenology from the Eastern Himalayan region.

## AUTHOR CONTRIBUTIONS

AD conceived, initiated, and acquired funding for the project. Field data was primarily collected by Khem T, Kumar T, TN, TB, NM with contributions from AD, RN, US, SS, AR, and NB. Data curation was done by AD, AR, SS, US, and SB. SB completed formal data analyses with contributions from AD and AR. AD and SB wrote the paper and all authors reviewed the paper.

## ACKNOWLEDGEMENTS

We thank the Arunachal Pradesh Forest Department for their support and research permits for this work, especially Pekyom Ringu, G.N. Sinha, Bharat Bhatt for their support in the past, and to N. Tam, Damodhar AT, Millo Tasser and Tajum Yomcha for their support. We are grateful to the Pakke Tiger Reserve management (Tana Tapi, SP Sinha, PB Rana, Rubu Tado, Kime Rambia and many frontline staff) for the various kinds of support and facilitating this work. We thank Rom Whitaker and the Agumbe Rainforest Research Station for providing the weather station and Shankar for technical support. We thank Kamal Sonar for maintaining the weather station. Arjun Rai, Sital Dako and several others have assisted with the tree phenology monitoring. We thank Veena Rai, Arjun Rai and Chaithra Gowda for assisting with data entry. Several grants for our long-term research and conservation program in the area enabled the continuation of the tree phenology monitoring – National Geographic Society, Disney Wildlife Conservation Fund, Wildlife Conservation Society, Whitley Fund for Nature, and the Serenity Trust.

## DATA AVAILABILITY STATEMENT

The data that support the findings of the study will be made openly available in Zenodo upon acceptance of the manuscript for publication. The corresponding DOI will be shared thereafter.

