## Supplemental Tables S1-S4, Figures S1-S3 for "Patterns of leaf, flower and fruit phenology and environmental relationships in a seasonal tropical forest in the Indian Eastern Himalaya"

SUPPLEMENTARY FILES

Table S1: Modal month of each phenophase in terms of the number of species and trees for years in which all months were sampled between January 2011 and February 2023 in Pakke Tiger Reserve. The month with the highest percentage of species or trees in the respective phenophase was designated as the modal month. Multiple months sharing the modal value are separated by “/”.

| Year | Percentage of species | | | | | | | Percentage of trees | | | | | | |
| --- | --- | --- | --- | --- | --- | --- | --- | --- | --- | --- | --- | --- | --- | --- |
|  |  |  |  | Ripe fruit (by seed dispersal mode) | | | |  |  |  | Ripe fruit (by seed dispersal mode) | | | |
|  | Leaf flush | Flower bud | Flower | Comm. | Bird | Mammal | Mech. | Leaf flush | Flower bud | Flower | Comm. | Bird | Mammal | Mech. |
| 2012 | May | April | May | October | April | October | February | June | March | May | May | May | October | February |
| 2013 | April | April | June | July | January/  March | June/July/  September | February/  March | April | April | May | June | May | October | September |
| 2014 | June | May | March | February | March | July | February/  September | May | May | March | April | March | October | March |
| 2016 | April | April | April | April | April | September/  October/  November | March | July | March | May | April | April | November | March |
| 2017 | April | March | March | January/  April | April | July | January | April | April | March | April | July | July | April |
| 2018 | April/  May | May | June | May | July | August | April | May | May | June | April | April | August | March |
| 2019 | May | April | April | November | May | November | February/  March | May | March/April | May | May | May | November | March |
| 2022 | April | April | April | June | June | September | January  /February/  March | April | April | March | April | April/  May | September | March |

^Comm.: Community-level	Mech.: Mechanically-dispersed^

Table S2: Model selection tables pertaining to the GAMLSS models used for modelling the relationships between the monthly proportion of trees in each phenophase or the fruiting intensity (monthly sum of the relative fruit abundance scores) and climatic variables. Model selection was carried out using the Akaike Information Criterion (AIC). AR(1) refers to the autoregressive term accounting for temporal autocorrelation by a period of 1 month.

| Model | AIC value | Delta AIC | AIC weight |
| --- | --- | --- | --- |
| **Leaf flush** | | | |
| ~mean minimum temperature+ AR(1) | -81.369 | 0 | 0.338 |
| ~daylength+ AR(1) | -81.086 | 0.282 | 0.293 |
| ~solar radiation+ AR(1) | -79.460 | 1.908 | 0.130 |
| ~solar radiation+ total rainfall+ AR(1) | -78.708 | 2.662 | 0.089 |
| ~total rainfall+ AR(1) | -77.871 | 3.498 | 0.059 |
| ~AR(1) | -77.322 | 4.047 | 0.045 |
| ~solar radiation+ proportion rainy days+AR(1) | -76.224 | 5.145 | 0.026 |
| ~proportion rainy days+ AR(1) | -75.573 | 5.796 | 0.019 |
| ~mean maximum temperature+ AR(1) | -70.558 | 10.811 | <0.001 |
| **Mature flower** | | | |
| ~mean minimum temperature+ solar radiation+ AR(1) | -317.186 | 0 | 0.473 |
| ~daylength+ AR(1) | -316.121 | 1.066 | 0.278 |
| ~mean minimum temperature+ AR(1) | -314.917 | 2.270 | 0.152 |
| ~total rainfall+ AR(1) | -312.989 | 4.197 | 0.058 |
| ~AR(1) | -310.523 | 6.663 | 0.017 |
| ~solar radiation+ total rainfall+ AR(1) | -309.561 | 7.626 | 0.010 |
| ~mean maximum temperature+ AR(1) | -308.057 | 9.130 | 0.005 |
| ~solar radiation+ AR(1) | -307.842 | 9.344 | 0.004 |
| ~proportion rainy days+ AR(1) | -306.587 | 10.599 | 0.002 |
| ~solar radiation+ proportion rainy days+ AR(1) | -303.409 | 13.778 | <0.001 |
| **Community-level fruiting** | | | |
| ~mean minimum temperature+ AR(1) | -400.889 | 0 | 0.447 |
| ~mean minimum temperature+ solar radiation+ AR(1) | -399.858 | 1.032 | 0.267 |
| ~daylength+ AR(1) | -398.517 | 2.373 | 0.135 |
| ~daylength+ solar radiation+ AR(1) | -395.765 | 5.124 | 0.034 |
| ~total rainfall+ AR(1) | -393.781 | 7.108 | 0.013 |
| ~AR(1) | -393.561 | 7.328 | 0.011 |
| ~solar radiation+ AR(1) | -392.576 | 8.313 | 0.007 |
| ~proportion rainy days+ AR(1) | -390.573 | 10.316 | <0.001 |
| ~solar radiation+ total rainfall+ AR(1) | -390.018 | 10.872 | <0.001 |
| ~mean maximum temperature+ AR(1) | -388.814 | 12.075 | <0.001 |
| ~proportion rainy days+ AR(1) | -388.775 | 12.114 | <0.001 |

Table S3: Estimates of beta coefficients and standard errors pertaining to climate variables used for modelling the proportion of trees in each phenological state. P values <0.05 indicate the significance of the cubic smoothed splines (denoted by cs()) associated with each climatic predictor variable. Estimates were derived from either the most parsimonious model or by model averaging across all models within 2 delta AIC of the best supported model.

| Parameter | Estimate (SE) | Z value | P value |
| --- | --- | --- | --- |
| Proportion of trees in leaf flush | | | |
| Intercept | 0.052 (0.025) | 2.105 | <0.05 |
| Sigma | -1.971 (0.076) | 25.813 | <0.0001 |
| cs(Mean minimum temperature) | 0.020 (0.018) | 1.127 | 0.260 |
| cs(daylength) | 0.049 (0.017) | 2.833 | <0.01 |
| cs(solar radiation) | 0.038 (0.016) | 2.418 | <0.05 |
| AR(1) | 0.720 (0.060) | 10.465 | <0.0001 |
| Proportion of trees in mature flower | | | |
| Intercept | 0.023 (0.007) | 3.204 | <0.01 |
| Sigma | -3.358 (0.080) | 41.772 | <0.0001 |
| cs(mean minimum temperature) | -0.011 (0.005) | 2.515 | <0.05 |
| cs(solar radiation) | 0.012 (0.004) | 2.719 | <0.01 |
| AR(1) | 0.626 (0.087) | 7.198 | <0.0001 |
| cs(daylength) | 0.002 (0.005) | -.410 | 0.682 |
| Percentage of trees in ripe fruit (community level) | | | |
| Intercept | 0.040 (0.005) | 8.264 | <0.0001 |
| Sigma | -3.834 (0.078) | 49.394 | <0.0001 |
| cs(mean minimum temperature) | 0.002 (0.003) | 0.693 | 0.488 |
| cs(solar radiation) | 0.006 (0.003) | 2.009 | <0.05 |
| AR(1) | 0.067 (0.096) | 0.700 | 0.484 |

Table S4: Cumulative AIC (Akaike Information Criterion) model weights corresponding to the cubic smoothed splines of each predictor obtained through GAMLSS modelling. The cs() term refers to the cubic smoothed splines of each climatic variable.

| Predictor | Leaf flush | Mature flower | Ripe fruit | Fruiting intensity |
| --- | --- | --- | --- | --- |
| Cs(Mean maximum temp) | <0.001 | <0.001 | <0.001 | 0.004 |
| Cs(Mean minimum temp) | 0.338 | 0.625 | 0.714 | 0.189 |
| Cs(Total rainfall) | 0.148 | 0.068 | 0.013 | 0.015 |
| Cs(Proportion rainy days) | 0.045 | 0.002 | <0.001 | 0.007 |
| Cs(Solar radiation) | 0.245 | 0.477 | 0.274 | 0.210 |
| Cs(Daylength) | 0.293 | 0.278 | 0.169 | 0.768 |

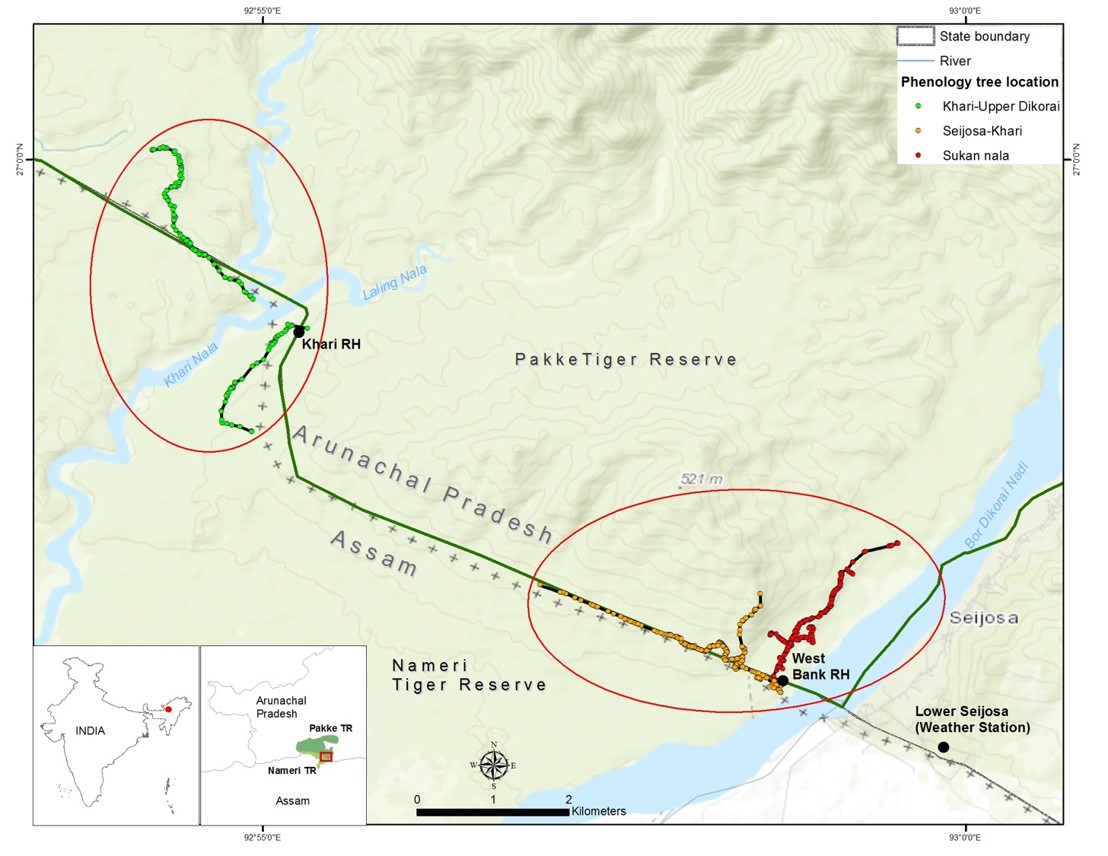

Fig. S1: Map of Pakke Tiger Reserve, Arunachal Pradesh, India, showing the locations of the major trails used for tree phenology monitoring.

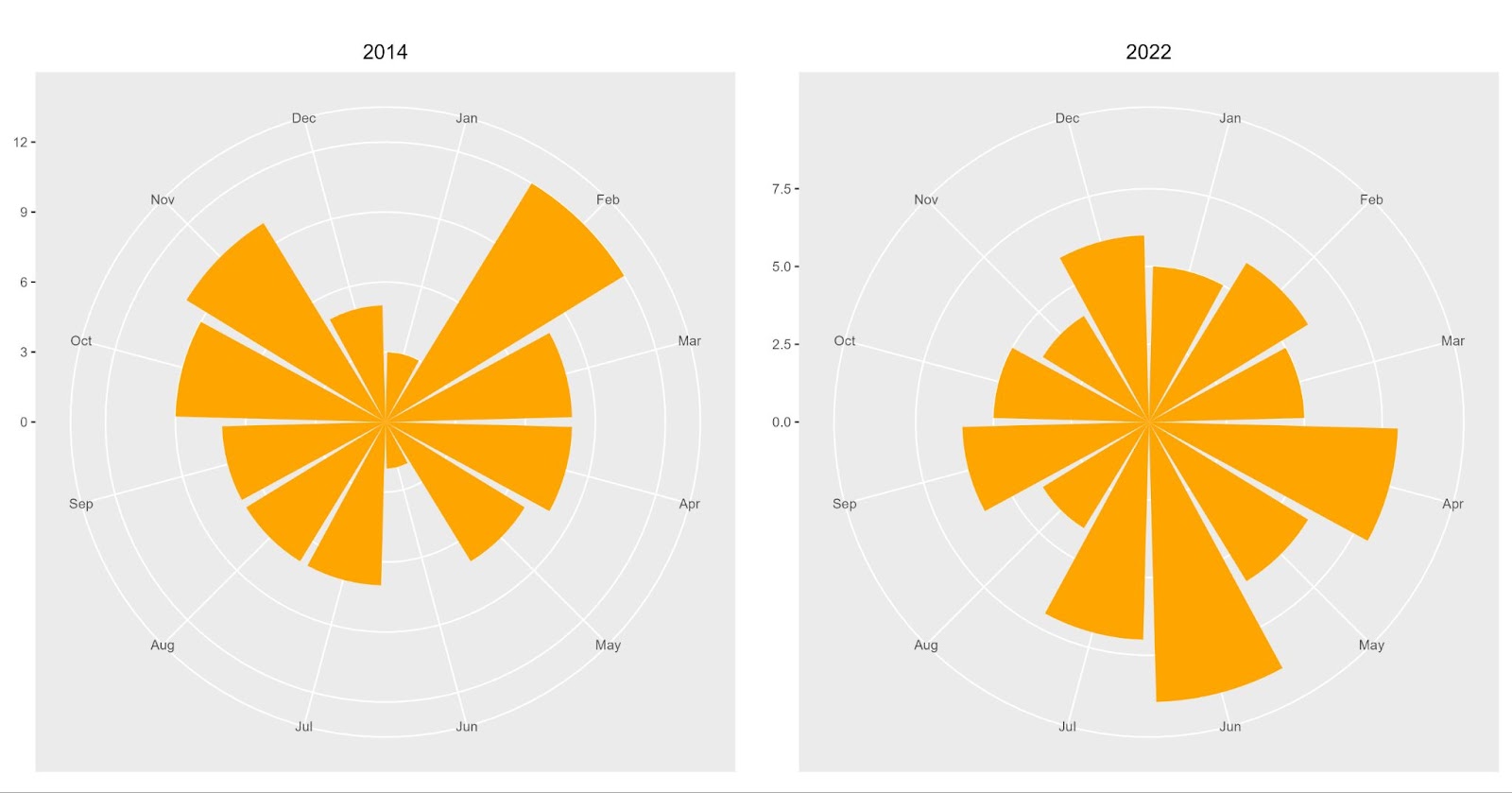

Fig. S2 a)  Circular histogram representing the distribution of the number of bird-dispersed species in the years 2014 and 2022 as histogram bars around the unit circle. The distribution is broadly bimodal for the year 2014 and uniform for the year 2022. The length of each bar represents the number of bird-dispersed species in fruit for each month.

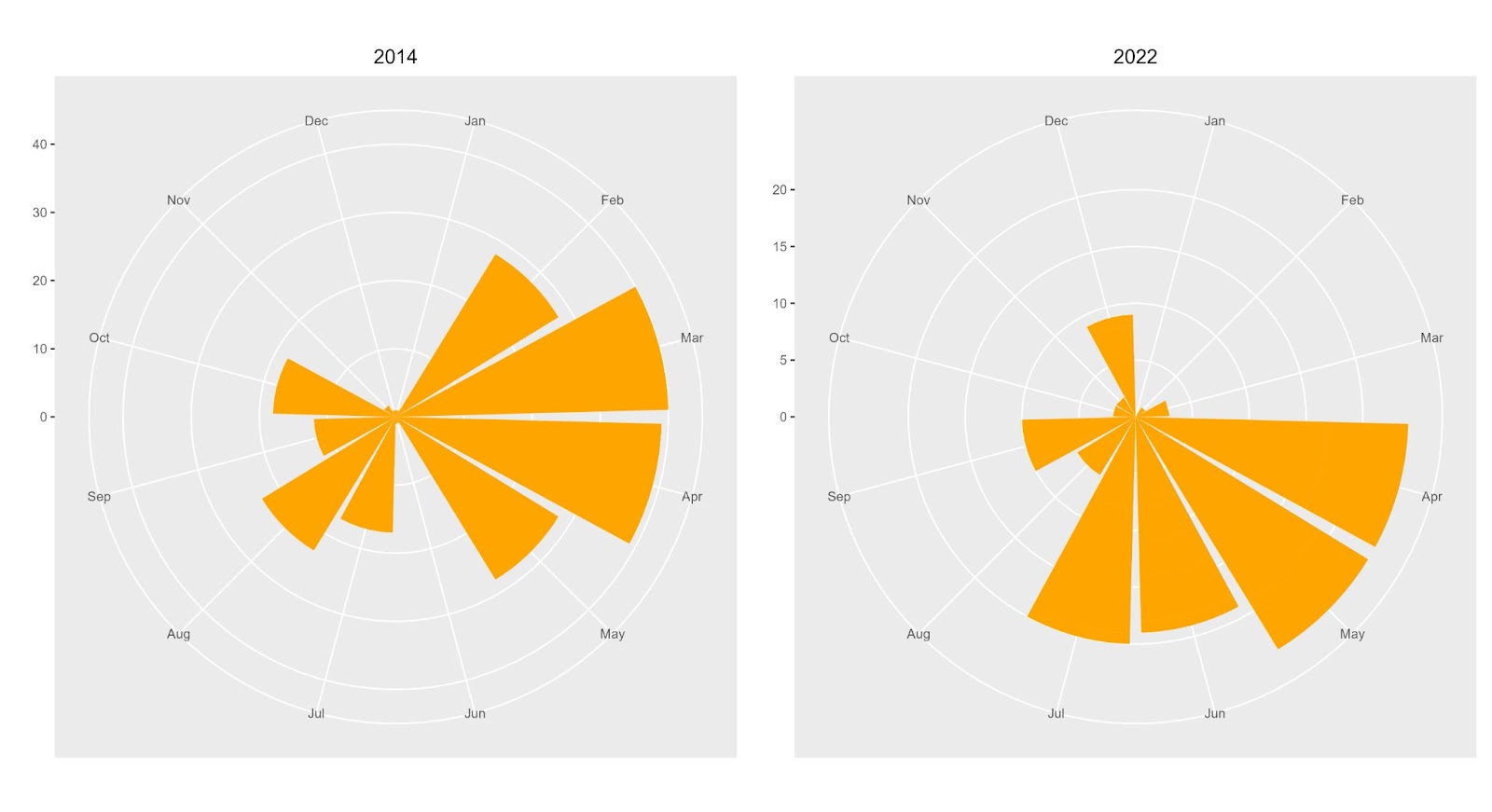

Fig. S2 b) Circular histogram representing the distribution of the number of bird-dispersed trees in the years 2014 and 2022 as histogram bars around the unit circle. The distribution was broadly bimodal in the year 2014 and unimodal in the year 2022. The length of each bar represents the number of bird-dispersed trees in fruit for each month.

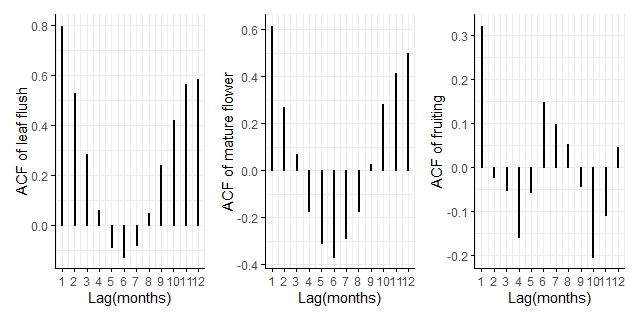

Fig.S3: Autocorrelation plots of the time series of the percentage of trees in leaf flush, mature flower and ripe fruit between January 2011 and February 2020.
